# Experimental Zika Virus Infection in a New World Monkey Model Reproduces Key Features of the Human Disease

**DOI:** 10.1101/102145

**Authors:** Charles Chiu, Jerome Bouquet, Tony Li, Shigeo Yagi, Claudia Sanchez San Martin, Manasi Tamhankar, Vida L. Hodara, Laura M. Parodi, Sneha Somasekar, Guixia Yu, Luis D. Giavedoni, Suzette Tardif, Jean Patterson

## Abstract

Human infections by Zika virus (ZIKV), a mosquito-borne flavivirus, are associated with a current widespread outbreak in the Americas, and have been associated with neurological complications and adverse fetal outcomes such as microcephaly in pregnant women. A suitable non-human primate model is urgently needed. To evaluate ZIKV infectivity, pathogenesis, and persistence, we inoculated 4 marmosets with ZIKV and followed them by clinical monitoring and serial sampling of body fluids for up to 11 weeks. We found that marmosets experimentally infected with ZIKV reproduced key features of the human disease, including (1) asymptomatic infection, (2) brief period of detectable virus in serum (<1 week), (3) detection in other body fluids (urine, saliva, semen, and stool) for at least 2 weeks following acute infection, and (4) persistence in lymph nodes, but not other tissues, at 1 month post-infection. ZIKV-positive saliva and serum samples, but not urine, were found to be infectious in cell culture. By day 6 post-inoculation, most marmosets exhibited detectable neutralizing antibody responses concurrent with activation of NK cell and B cell subsets and an increase in circulating cytokines associated with type II interferon signaling, Transcriptome profiling revealed enrichment of immune responses to active viral infection, with up-regulation of both type I and II interferon signaling pathways, and uncovered potential host biomarkers. These results suggest that a New World monkey model of acute ZIKV infection mimics the human disease, and is likely to be useful for testing of drug and vaccine candidates.

## IMPORTANCE

A monkey model of Zika virus (ZIKV) infection is urgently needed to better understand how the virus is transmitted and how it causes disease, especially given its proven association with fetal brain defects in pregnant women and acute neurological illness. Here we experimentally infected 4 marmosets, which are New World monkeys, with ZIKV and monitored them clinically with sampling of various body fluids and tissues for nearly 3 months. We show that the course of acute infection with ZIKV in marmosets resembles the human illness in many respects, including (1) lack of apparent clinical symptoms in most cases, (2) persistence of the virus in other body fluids for longer periods of time than blood, and (3) generation of a (neutralizing antibodies as well as an antiviral immunological host response. Importantly, ZIKV-infected saliva samples (in addition to blood) were found to be infectious, suggesting potential transmission of the virus by the oral route. Our data indicate that marmosets may be a promising animal model for ZIKV infection to test new drug and vaccine candidates.

## INTRODUCTION

Zika virus (ZIKV) is an infectious RNA flavivirus primarily transmitted to humans by the bites of *Aedes* spp. mosquitoes (1). An outbreak of ZIKV began in Brazil in early 2015 and has since spread throughout South America, Central America, and the Caribbean, with autochthonous cases now being reported in the United States (Miami, Florida, and Texas). The rapid emergence of ZIKV in the Western Hemisphere is of particular concern given the proven association of viral infection with devastating fetal outcomes in pregnant women, including miscarriage and microcephaly (2). Although the majority of ZIKV-infected individuals (~80%) are asymptomatic (3), patients can present with a self-resolving acute illness consisting of fever, conjunctivitis, rash, and joint pain. Rarely, ZIKV has also been associated with rare neurological complications such as meningoencephalitis (4) and Guillain-Barre syndrome (5).

Although the primary mode of ZIKV transmission is via mosquito bite, it has also been shown that the virus has the capacity for sexual transmission (6). Following an acute infectious episode, the virus can reside in semen for at least 3 months (7). The virus has also been detected for at least 2 weeks after symptom onset in saliva and urine samples from acutely infected individuals (8), although it is unknown whether the these sampled body fluids were infectious. ZIKV transmission by blood transfusion from an infected donor has also been reported (9).

To date, there have been several published mouse models of ZIKV infection; however, these have focused on studying ZIKV-associated complications in pregnant females such as fetal microcephaly (10,-13), and require the use of immunodeficient animals with defects in interferon-related signaling pathways, likely due to absence of STAT2 cytokine inhibition of ZIKV in mice (14). A viable nonhuman primate (NHP) model may thus better reflect the biology and pathogenesis of ZIKV in acute human infections. Investigations with NHP can also enable serial sampling and analyses of body fluids (e.g. urine, saliva, feces, and semen) that are impractical with rodent models.

Rhesus macaque (*Macaca mulatta*) models of ZIKV infection are currently in development (15). However, there are compelling reasons to consider the common marmoset (*Callithrix jacchus*), a New World monkey, as a useful alternative candidate model for ZIKV investigation. Common marmosets are known to have a high susceptibility to infection by a variety of pathogenic outbreak agents (16), including Ebola virus (17), Lassa virus, (18), and titi monkey adenovirus − a virus found to be associated with cross-species transmission to both monkeys and humans (19). Related flaviviruses to ZIKV, including dengue virus (DENV) and West Nile virus (WNV) are known to cause productive infections in marmosets (20, 21). In addition, the recent detection of ZIKV in serum or saliva from a high percentage of wild marmosets from Brazil (26.7%, 4 of 15 animals tested) (22) suggests that marmosets can harbor ZIKV, and thus may constitute a reservoir for maintaining the virus in the wild.

Here we present a marmoset model of acute ZIKV infection generated by inoculating 4 animals with the prototype 1947 Uganda strain of ZIKV (23), followed by clinical monitoring and serial sampling for nearly 3 months. We sought to evaluate ZIKV infectivity, pathogenesis, persistence in infected body fluids and potential transmission risk, and production of neutralizing antibodies. The host response to acute ZIKV infection was also investigated by lymphocyte phenotyping, cytokine analyses and global transcriptome profiling of blood from experimentally infected animals.

## METHODS

### Animal Ethics Statement

All animal studies were conducted at the Southwest National Primate Research Center (SNPRC), Texas Biomedical Research Institute (TBRI); molecular, viral, and transcriptome analyses of marmoset body fluids and tissues were conducted at University of California, San Francisco (UCSF). TBRI is accredited by the Association for Assessment and Accreditation of Laboratory Animal Care (AAALAC) International and operates in accordance with the NIH and U.S. Department of Agriculture guidelines and the Animal Welfare Act. The Institutional Animal Care and Use Committee (IACUC) and the Institutional Biohazards Committee (IBC) of the TBRI approved all marmoset protocols related to this study.

Marmosets were kept healthy and well nourished with strict feeding protocols and close monitoring of their health status prior to the start of the study and during the entire study period. One week before inoculation, animals were transferred to the biosafety level-2 facility at the SNPRC and housed individually in cages specifically developed for marmoset work. As they are social animals in the wild, all marmosets had auditory, visual, and olfactory access to each other throughout the study. Marmosets were sedated and humanely euthanized by administration of a sodium pentobarbital solution by a licensed veterinarian at the TBRI.

### ZIKV propagation in cell culture

Vero cells were inoculated with the 1947 Uganda strain of ZIKV (passaged 147X in mouse brain and 3X in Vero cells) in the African lineage, which has been maintained at the Viral and Rickettsial Disease Laboratory (VRDL) branch of the California Department of Public Health. Viral supernatants for cell culture passaging and the generation of infectious stocks were subjected to 3 freeze-thaw cycles and clarified by centrifugation for 10 min × 4000 *g*. After cells achieved 80–90% confluency, cell culture media were changed to maintenance media with 2% FBS and were inoculated with 100 μL of passaged viral supernatant. Viral replication was monitored over 14 days by visual inspection under light microscopy for cytopathic effect (CPE). Infectious viral titers were quantified using an end-point dilution assay.

### Experimental ZIKV infection of marmosets

Four healthy adult male marmosets, averaging 2.1 years of age (range: 2.0 – 2.3 years) and 391.7 g (range: 332 – 453 g), were inoculated intramuscularly with 1x10^5^ pfu/mL of the 1947 Uganda African lineage of ZIKV. Samples from an additional 4 male marmosets were used as matched controls and to establish a transcriptomic baseline. Study marmosets were screened for ZIKV antibody by neutralization and were found to be negative.

Animals were monitored daily for signs of clinical illness, with generalized sickness defined as a score of >4 (**Supplementary Table 1**). Specific monitoring was conducted for signs associated with ZIKV infection in humans, including rash, anorexia, conjunctivitis, diarrhea, malaise, and postural abnormalities associated with joint or head pain. Clinical samples were collected from restrained, unsedated animals at predetermined time intervals. Animals were restrained for less than 10 minutes in a device specifically designed for short-term restraint of marmosets for sample collection purposes. Blood samples were collected via venipuncture on Days 1, 3, 6, 9 and 28; voided urine and feces were collected on days 3, 5, 7, 9, 11, and 13; saliva was collected on days 3, 6, 9, and 14 by allowing the subjects to chew on a sterile cotton swab; and semen samples were collected on days 9, 14, 28 by vibratory stimulation of the penis, using a modified FertiCare^TN^ medical vibrator unit (Multicept A/S, Denmark) (**Fig. 1**). Whole blood was collected in tubes containing RNA stabilization media (Biomatrica, Inc.) for transcriptome analysis. At day 28, 2 of the 4 animals were randomly selected to be euthanized. The remaining two inoculated marmosets were observed for an additional 7 weeks, with clinical samples collected at weeks 7, 10 and/or 11 to evaluate long-term ZIKV persistence in body fluids.

**Figure 1.**
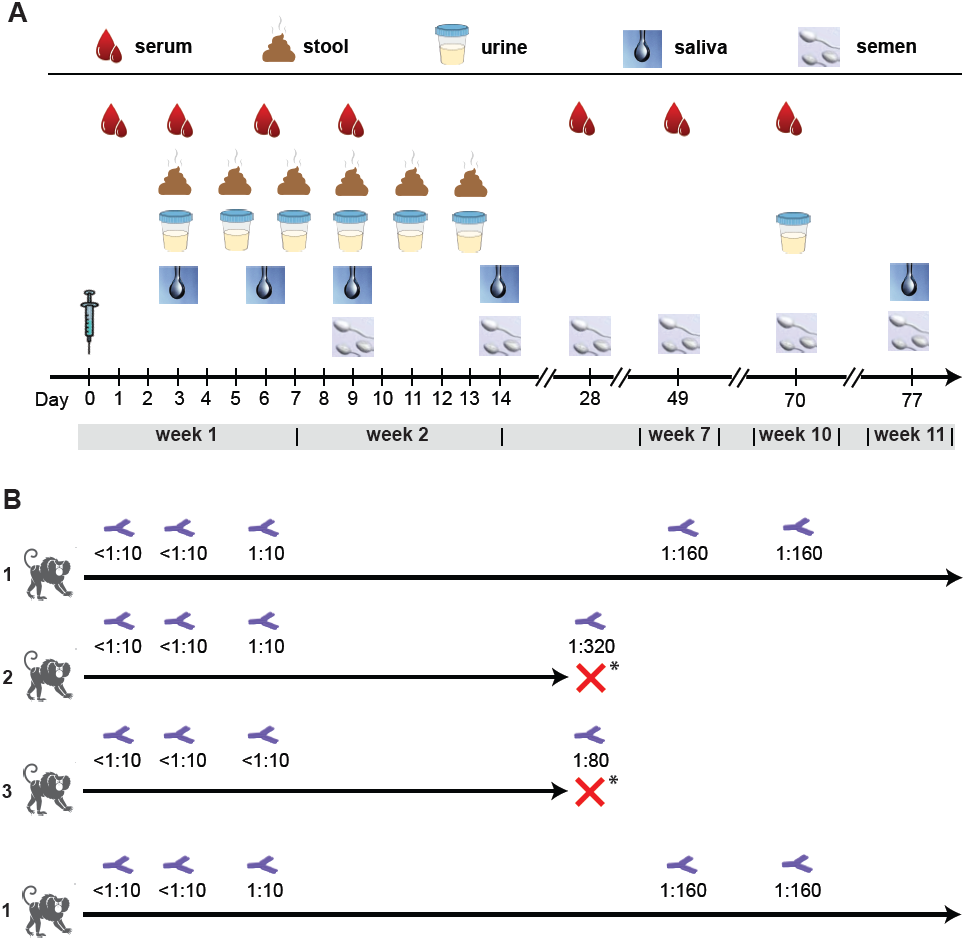
**Study design and neutralizing antibody testing**. (A) After intramuscular inoculation of ZIKV at day 0, samples (serum, stool, urine, saliva, and semen) are collected at pre-established time points. (B) Longitudinally collected serum samples from inoculated marmosets were tested at pre-established time points for ZIKV-specific neutralizing antibodies using a PRNT (plaque reduction neutralization test). Two of the 4 marmosets were sacrificed at day 28 to assess viral persistence in tissues.

### Measurement of ZIKV titers from clinical samples by qRT-PCR

The course of infection was monitored by determination of ZIKV titers in serum, urine, saliva, stool, semen (collected in a conical tube at the time of ejaculation), and semen swabs (semen swabbed off of the penis and surrounding tissues immediately following ejaculation). Viral titers were calculated by generation of a standard curve, followed by quantitative RT-PCR testing using primers targeting the envelope gene (ZIKV-1086/ZIKV-1162), as previously described (24).

### ZIKV serological analysis by antibody neutralization

Plaque-reduction neutralization testing (PRNT) on longitudinally collected marmoset sera was performed by the California Department of Public Health. The protocol was similar to that used by the US CDC for confirmatory ZIKV testing in patients (25). Briefly, 100 plaque forming units (PFU) of ZIKV (Uganda strain) were mixed with equal volumes of serial 2-fold dilutions of inactivated marmoset sera and incubated for 1 hr at 36°C, followed by inoculation and adsorbing to a monolayer culture of Vero cells for 1 hr at 36°C. After addition of 3 mL of 1% agar in Eagle’s Minimal Essentail Medium (MEM), plates were placed in a 36°C, 5% CO_2_ incubator x 3 days, followed by addition of 3 mL of 1% agar and 0.004% neutral red in Eagle’s MEM and another 1-2 days of incubation until plaques were formed. An 80% reduction of the number of plaques compared to positive control wells inoculated with virus-diluent mixtures was considered neutralization, with serum titers reported as the highest dilution exhibiting ≥80% reduction.

### Histology of ZIKV-infected marmosets

Samples of aseptically removed tissues were fixed in 10% neutral buffered formalin and embedded in paraffin for histology. Paraffin-embedded tissues were cut in 5 μm sections, deparaffinized, and stained with hematoxylin and eosin (H&E) prior to visualization by light microscopy. Additional samples were freshly frozen in liquid nitrogen and kept stored in a −80°C freezer until analyzed. Two board-certified veterinary pathologists independently evaluated the histologic sections.

### Lymphocyte phenotyping

Phenotypic characterization of marmoset PBMCs was performed by multicolor flow cytometry using direct immunofluorescence. Aliquots of 100 μl of EDTA whole blood were directly incubated with antibodies for 20 minutes at room temperature; red blood cells were lysed with ACK, and cells were then washed twice with PBS and fixed with 1.6% methanol-free formaldehyde before analysis in a CyAn LDP flow cytometer (Beckman-Coulter). The antibodies used for this analysis were conjugated to fluorescein isothiocyanate (FITC), Phycoerythrin (PE), Peridinin-chlorophyll-cyanin 5.5 (PerCP-Cy5.5), Phycoerythrin-cyanin 5.1 (PC5), Phycoerythrin-cyanin 7 (PC7), Pacific Blue, BD Horizon V500, Allophycocyanin (APC) or Alexa Fluor 700. Antibodies included in this study were: CD3 (clone SP34.2), CD4 (clone L200) and HLA-DR (clone G46.6/L243) from BD-Biosciences; CD14 (clone 322A-1 (My4), CD159a (NKG2A; clone Z199), CD20 (clone H299(B1)), CD335 (NKp46; clone BAB281) and CD337 (NKp30; clone Z25) from Beckman-Coulter; CD16 (clone 3G8), CD8 (clone HIT8a), CD86 (clone IT2.2) from Biolegend; and CD159c (NKG2C; clone 134522) from R&D Systems.

For analyses, lymphocytes were gated based on their characteristic forward and side scatter pattern, followed by T-cell selection using a second gate on the CD3-positive population. Thus, CD8 T cells were defined as CD8^+^/CD3^+^ and CD4 T cells as CD4^+^/CD3^+^. Natural Killer cells (NK) were defined as CD3^-^/CD20^-^/CD14^-^ and analyzed by the expression of NK cell markers CD16^+^, CD8, NKG2A, NKG2C, NKp30 and NKp46. B cells were defined as CD20^+^/CD3^-^/CD14^-^.

### Multiplex cytokine analysis of plasma

Plasma samples were analyzed for marmoset cytokines and chemokines on the Luminex 100 system (Luminex) using established protocols for New World primates (26). The assay included evaluation of the following 21 analytes: GRO-α (CXCL1), interferon alpha (IFN-α), IFN-γ, interleukin-1 beta (IL-1β), IL-1 receptor antagonist (IL-1RA), IL-4, IL-8, IL-10, IL-12 p70, IL-15, IL-18, IL-22, monocyte chemoattractant protein 1 (MCP-1, CCL2), macrophage migration inhibitory factor (MIF), monokine induced by gamma interferon (MIG, CXCL9), macrophage inflammatory protein 1-alpha (MIP-1α, CCL3), MIP-1β (CCL4), regulated on activation, normal T cell expressed and secreted (RANTES, CCL5), tumor necrosis factor-alpha (TNF-α), soluble CD40 ligand (sCD40L), soluble intercellular adhesion molecule 1 (sICAM-1), and vascular endothelial growth factor A (VEGF-A).

### Transcriptome analysis

Four age-/sex-matched healthy marmosets were used as controls for the transcriptome analysis. All marmosets studied here were from a single colony, thus increasing genetic similarities and decreasing environmental bias. Technical bias in the whole transcriptome analysis was not observed by PCA (**Supplementary Fig. 1**)

Four hundred microliters of blood were drawn directly into RNAgard tubes (Biomatrica) for immediate RNA stabilization of intracellular RNA at collection. Total RNA was extracted using the Biomatrica Blood RNA Purification Kit (Biomatrica). The Ovation Human Blood RNA-Seq Kit (Nugen) was used to generate RNA-seq libraries according to the manufacturer's protocol. Libraries were sequenced as 100 base pair (bp) paired-end runs on a HiSeq 2500 instrument (Illumina).

Paired-end reads were mapped to the marmoset genome (Callithrix jacchus Ensembl version 3.2.1), using STAR 2.5 (27), and gene and transcript normalized counts were calculated by HTSeq version 0.6.0 (28). Differential expression of genes was calculated using linear modeling using the Bioconductor EdgeR software package version 3.12.2 (29) implemented in the R programming language. Genes were considered to be differentially expressed when their fold change was > ± 2, *p*-value < 0.05, and adjusted *p*-value (or false discovery rate, FDR) < 0.1%. Pathway and network analyses of the transcriptome data were performed using Ingenuity Pathway Analysis (IPA) software (Qiagen).

Marmoset transcriptome data has been submitted to the public National Center for Biotechnology Information (NCBI) Gene Expression Omnibus (GEO) repository (accession numbers pending).

## RESULTS

### Experimental infection of 4 male marmosets with ZIKV

To investigate ZIKV infectivity and potential pathogenesis, we inoculated 4 healthy male marmosets intramuscularly with 0.1 mL of 10^5^ plaque-forming units (PFUs) of the 1947 Uganda prototype strain of ZIKV. The inoculation dose was chosen to be physiologic, comparable to the highest observed serum titers in patients with acute ZIKV infection (30). Marmosets remained largely asymptomatic during the entire study period, with the exception of one animal who exhibited drowsiness 2 days post-inoculation and had lost 7% of its body weight by day 5. However, this animal subsequently appeared alert and active and ate normally. No animal ever displayed a clinical score of >=4, indicative of acute sickness, at any time during the study (**Supplementary Table 1**). Specifically, none of the other inoculated animals displayed anorexia, activity changes, or weight loss, and no subjects had fever, rash, conjunctivitis, diarrhea, or postural abnormalities suggestive of joint and/or muscle pain.

### ZIKV titers in body fluids from experimentally infected marmosets

Serum, saliva, and urine samples were collected longitudinally from sedated animals at predetermined intervals (see Methods) for up to 14 days following inoculation and levels of ZIKV measured by quantitative ZIKV RT-PCR (**Fig. 1A; Fig. 2; Supplementary Table 2**). A rapid rise and fall in viral titer, beginning at day 1 and returning to zero within 7-9 days, was observed in sera from inoculated animals (**Fig. 1A**). Peak viremia was >10^5^ copies/mL in all 4 animals at day 3 post-inoculation. In contrast to serum, viral titers in urine and saliva persisted for longer periods of time, with peak viral titers comparable to those observed in serum. Notably, at the end of the ~14day collection period, 3 of 4 animals (75%) and 2 of 4 animals (50%) were still shedding virus in the urine and saliva, respectively. Virus was also detected in the feces of inoculated animals from beginning on day 5, albeit at much lower titers, and one animal (25%) continued to shed virus at day 13. The virus was also sporadically detected in semen and semen swabs in some, but not all, animals at low titers during the first 2 weeks following inoculation.

**Figure 2.**
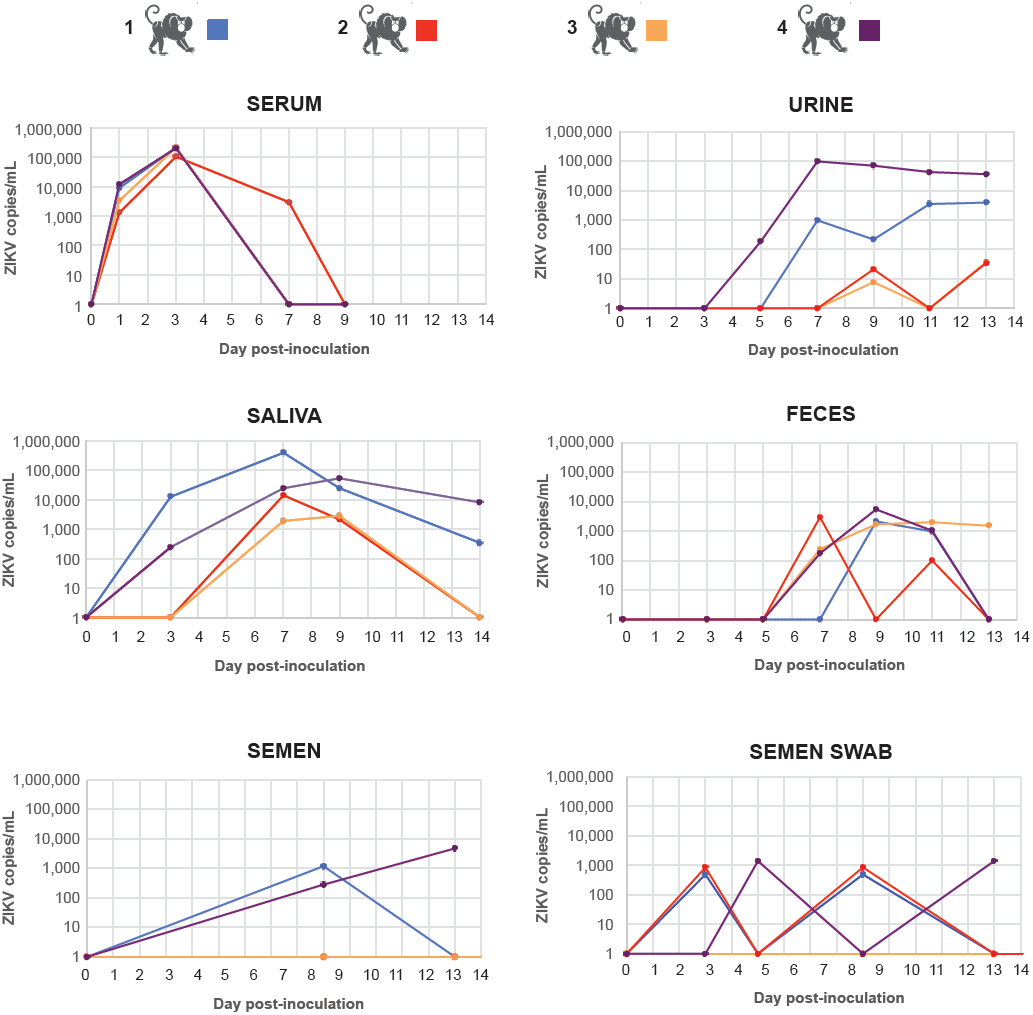
**Viral loads in body fluids after acute ZIKV infection**. The ZIKV load in copies per milliliter is plotted according to day post-inoculation. The line graph corresponding to each marmoset is displayed in a separate color.

A ZIKV antibody neutralization assay by plaque reduction neutralization testing (PRNT), validated at the California Department of Public Health, was used to screen experimentally infected marmosets for the development of neutralizing antibodies (Abs) to ZIKV. Importantly, all 4 marmosets had negative pre-inoculation Ab titers of <1:10. ZIKV neutralizing antibody responses were detected in all 4 animals at a titer of 1:10 by day 6 post-inoculation, and positive titers ranging from 1:80 – 1:320 after week 4 (**Fig. 1B**).

### Gross pathology and histology in ZIKV-infected marmosets

At 4 weeks post-inoculation, 2 of the 4 marmosets were randomly selected to be humanely euthanized for analysis of ZIKV pathology and persistence in tissues (**Fig. 1B**). No significant gross pathological lesions were observed in any of the post-mortem tissues. Salient histologic findings include mild-moderate nephropathy and vacuolization in hepatocytes associated with glycogen storage. Both of these histologic findings are common in healthy marmosets from this colony.

### ZIKV persistence in tissues and body fluids from experimentally infected marmosets

From the 2 sacrificed marmosets, no virus was detectable by qRT-PCR from necropsy conjunctival, kidney, heart, liver, lung, midbrain, prostate, salivary gland, spinal cord, or testicular tissue (**Supplementary Table 2**). One of the 2 marmosets had detectable virus (3,680 copies/mg) in a lymph node. To evaluate long-term persistence in body fluids, we also collected semen (6 and 10 weeks), semen swab (day 42), serum (6 and 10 weeks), urine (10 weeks), and saliva (11 weeks) from the 2 remaining marmosets. None of the samples collected after 6 weeks were positive for ZIKV by qRT-PCR.

### Viral infectivity from serum, saliva, and urine

Next, we sought to determine whether detected virus in body fluid compartments (e.g. serum, saliva, and urine) was infectious. We inoculated Vero cells with available ZIKV RT-PCR positive body fluids from infected marmosets at titers of 3.4x10^2^ to 1.9x10^5^ copies (**Table 1**). Viral cytopathic effect was observed after inoculation of all 4 of 4 day 3 serum samples (each collected from an individual marmoset), and 4 of 4 day 7 saliva samples, but not from urine samples or day 14 semen samples.

**Table 1.**
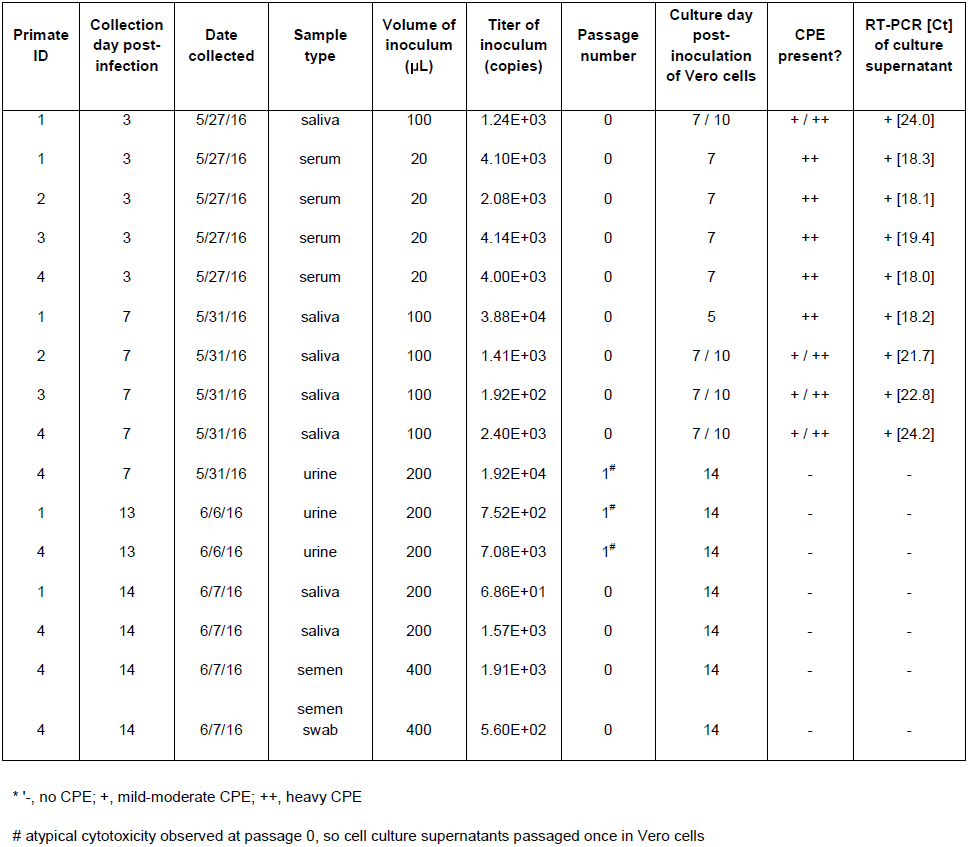
**Cell culture of ZIKV from infected body fluids**. Abbreviations: CPE, cytopathic effect; RT-PCR, reverse-transcriptase polymerase chain reaction; Ct, cycle threshold.

### Cytokine and lymphocyte analyses

Flow cytometry analysis of circulating lymphocytes in ZIKV-infected marmosets showed no major changes for most of the lymphocyte subsets that were studied, including levels of T cells or CD8 T cells (**Fig. 3A**). However, we did observe an increase in the population of NKG2A+ NK cells, which peaked by days 7-9 post-infection and returned to pre-infection levels by day 28 post-infection. There were also detectable increases in the levels of the NK activation markers NKp30, NK p46 and XX (data not shown). Interestingly, there was also a continuous up-regulation of the activation markers CD86 and HLA-DR on B cells during this acute period, returning to pre-infection levels by day 28 post-infection.

**Figure 3.**
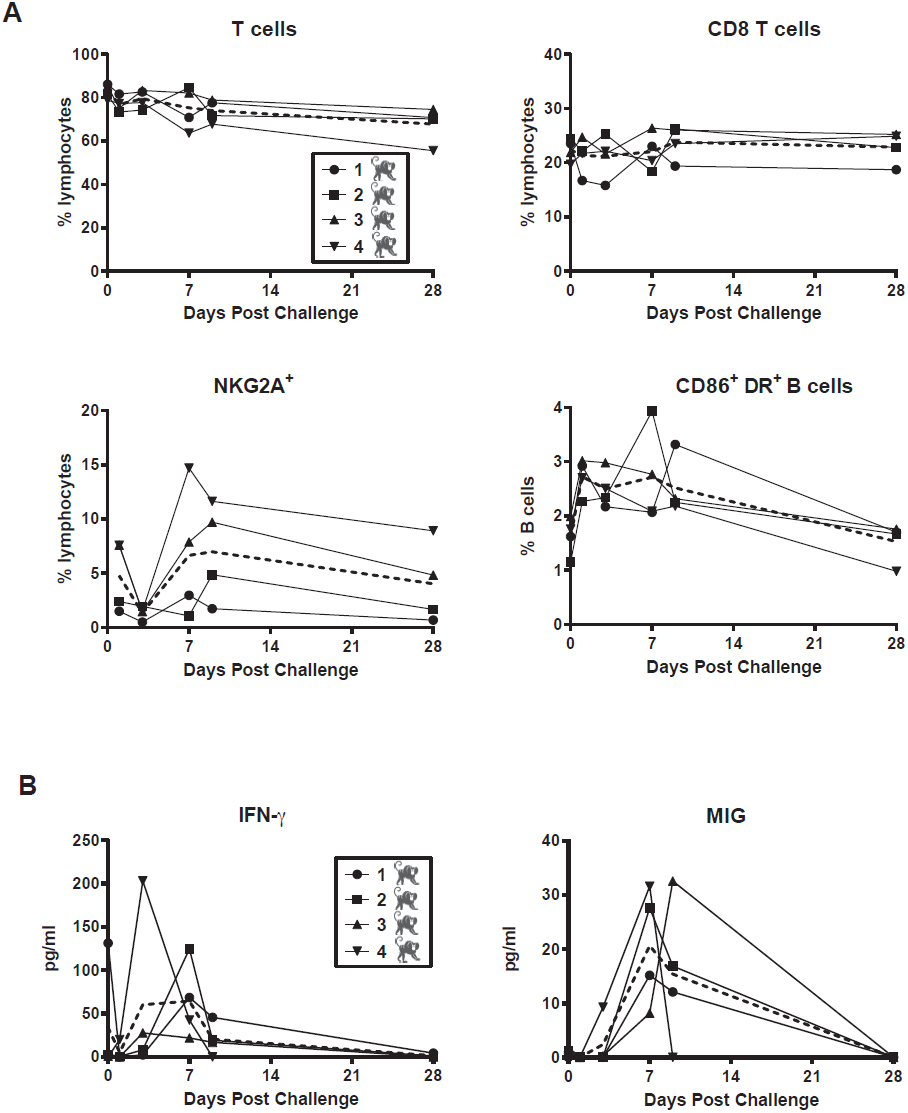
**Changes in lymphocyte subsets and circulating cytokines after acute ZIKV infection**. (A) Polychromatic flow cytometry was used to identify T cells (upper left), CD8 T cells (upper right), NKG2A+ NK cells (lower left), and CD20+ B cells expressing activation markers CD86 and HLA-DR (lower right). (B) Increases in protein expression of interferon-gamma (IFN-γ) and monokine induced by IFN-γ (MIG) were detected using a Luminex assay.

In parallel, we determined the plasma levels of 21 cytokines and chemokines with a validated Luminex assay (26). The majority of these molecules were either below the limit of detection of the assay or did not change in a significant way after challenge with ZIKV (**Supplementary Table 3**). However, there was an increase over time in circulating IFN-γ and MIG (CXCL9, a monokine induced by IFN-γ), both members of the type II interferon signaling pathway, which peaked between days 3 and 9 post-infection, and returned to basal levels by day 28 post-infection (**Fig. 3B**). In contrast, circulating levels of IFN-α, representative of the antiviral type I interferon response, were always below the limit of detection (**Supplementary Table 3**).

### Whole transcriptome data analysis

The 4 asymptomatic ZIKV-infected male marmosets were sampled for whole blood transcriptome analysis at days 1, 3, 7, and 9 after post-infection, and were compared to 4 healthy uninfected male marmosets as controls. Two of the marmosets were followed up for 24 days post-infection and the remaining two marmosets were continually followed up for 42 and 64 days post-infection. The average sequencing depth was 25.0M reads per sample (±13.9M reads) (**Supplementary Fig. 2**). STAR/Cufflinks detected an average of 51.2% (±9.9%) of all Ensembl isoforms in each sample.

We examined whether there were any differentially expressed genes (DEGs) between the Zika-infected marmosets at 1, 3, 7, 9, 42, and 64 days after post-infection and controls from their whole blood samples. For the first day post-infection (D1), 3 DEGs were found comparing Zika-infected against uninfected marmosets (**Table 2**). Three days post-infection (D3), 20 DEGs were found between the Zika-infected marmosets and controls, with 90% (n=18) up-regulated and 10% down-regulated (n=2). At seven days post-infection (D7), the difference in gene expression increased relative to the controls, with 43 DEGs found, 95% (n=41) up-regulated and 5% (n=2) down-regulated. By nine days post-infection (D9), 1,049 DEGs were found, with 67% (n=706) up-regulated and 33% (n=343) down-regulated. Two animals were sacrificed after day 9, and only 2 marmosets remained for follow up at day 42 and 64 post-infection, with 12 and 20 DEGs found respectively, but significance is uncertain given the low number of replicates. Twelve DEGs, all up-regulated, were shared between D3, D7, and D9 time points (**Supplementary Table 4**). One DEG, U3, a small nucleolar RNA, was shared between all time point.

**Table 2.** **Number of differentially expressed genes (DEG) between ZIKV-infected and uninfected marmosets by day post-inoculation**.

| **Comparison** | **Total DEGs** | **Up-regulated** | **Down-regulated** |
| --- | --- | --- | --- |
| Day 1 versus uninfected | 3 | 1 | 2 |
| Day 3 versus uninfected | 20 | 18 | 2 |
| Day 7 versus uninfected | 43 | 41 | 2 |
| Day 9 versus uninfected | 1049 | 706 | 343 |
| Day 42 versus uninfected | 12 | 5 | 7 |
| Day 64 versus uninfected | 20 | 9 | 11 |
| Zika (all time points) versus uninfected | 6 | 3 | 3 |

Gene ontology analysis revealed 12 DEGs shared between days 3, 7, and 9 comparison of ZIKV-infected versus uninfected marmosets (**Supplementary Table 4**). Among the DEGs, there was an enrichment of terms related to the defense response to virus (GO:0051607), innate immune response (GO:0045087), and negative regulation of viral genome replication (GO:0045071). Notably, 6 of the 12 DEGs (MX1, MX2, ISG15, OAS2, OAS3, and GP2) were members of the type I interferon signaling pathway, whereas 1 DEG (GBP1) was a member of the type II interferon signaling pathway.

Canonical pathway analysis showed that the interferon signaling pathway, which regulates host resistance against viral infections, was the only pathway significantly up-regulated at all blood sampling time points: 3 (n=20 pathways), 7 (n=22 pathways), and 9 (n=53 pathways) post-infection (**Fig. 4**). Type I interferon pathway was activated at day 3, 7 and 9, and type II interferon pathway was activated at day 9 (**Supplementary Fig. 3**). Pathways related to cell activation: eIF2 signaling, actin-based motility by Rho family, and RhoA signaling pathways, were found to be significantly up-regulated only at D9. Pathway analysis at days 42 and 64 post-infection was attempted, but no pathway with significant up- or down-regulation could be predicted (data not shown).

**Figure 4.**
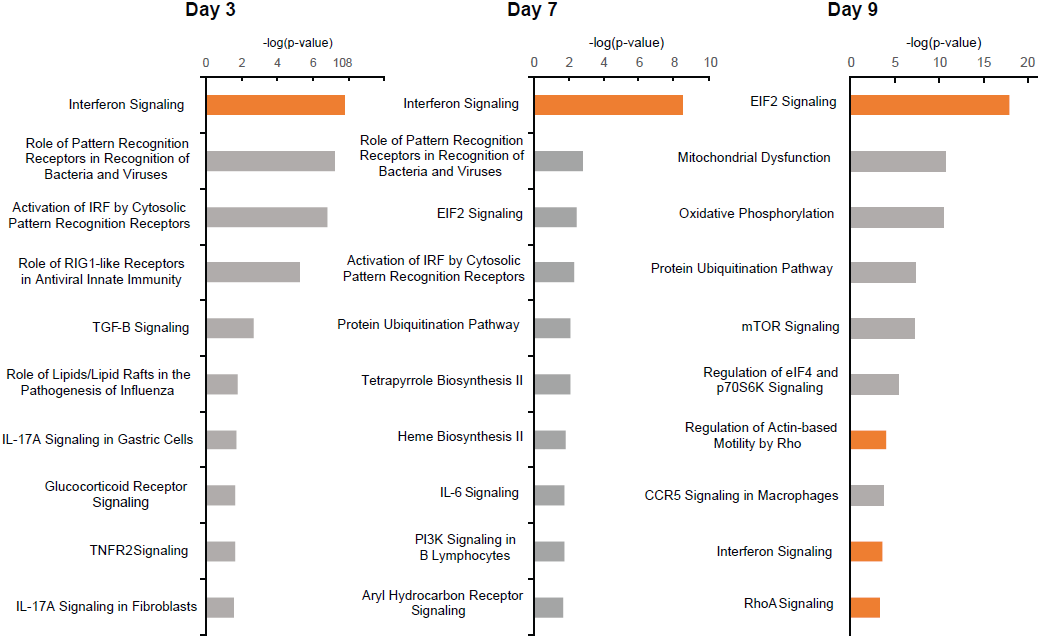
**Top 10 canonical pathways associated with acute ZIKV infection by transcriptome profiling**. Shown are the top 10 pathways at days 3, 7, and 9 post-inoculation, ranked by the negative log of the P-value of the enrichment score. The color scheme is based on Z-scores, with activation in orange and undetermined directionality in gray.

## DISCUSSION

In this study, we found that marmosets inoculated with ZIKV did not develop signs of clinical illness, mimicking the approximately 80% of human infections that are asymptomatic (3). Also resembling human infections are (1) the brief period of viremia (< 1 week), (2) persistent elevated titers in saliva and urine for at least 2 weeks following infection, and (3) sporadic detection in semen and stool. Analysis of differential gene expression in ZIKV-infected marmosets revealed specific enrichment of immune responses to viral infection, with antiviral type I interferon signaling as the only pathway significantly up-regulated at all 3 sampled time points (days 3, 7, and 9 days post-infection), However, cytokines associated with type II interferon signaling (IFN-γ and MIG), but not type I, were detected in sera, concurrent with systemic activation of NK cell and B cells subsets and robust generation of anti-ZIKV neutralizing antibodies. Importantly, we report that saliva and serum, but not urine, from ZIKV-inoculated animals were found to be infectious. Taken together, these results indicate that the ZIKV marmoset model mimics important aspects of the human disease.

Peak ZIKV titers in saliva and urine were comparable to those observed in serum and persisted for at least 2 weeks. This is consistent to what has been previously documented in humans, and incorporation of these additional sample types is now part of many diagnostic and public health surveillance efforts. It is also notable that ZIKV persistence in saliva and urine, unlike in blood, was not uniform, despite the fact that the marmosets were derived from a closed colony suggesting a population that was relatively homogeneous genetically. Thus, host and perhaps environmental factors may likely play an important role in determining the degree of ZIKV shedding in a particular individual, as has been shown for certain patients with acute ZIKV infection (31, 32)

In 1 of 2 marmosets that were euthanized at 28 days, we also found evidence of ZIKV persistence in lymph node tissue, but not other examined tissues. Lymphoid tissues are known “sanctuary sites” for viral infections such as HIV-1 (33), and, interestingly, acute ZIKV infection associated with lymphadenopathy has been described (34). As samples were collected infrequently past the 2-week post-inoculation period, repeat studies in marmosets with more regular sampling will be needed to precisely assess persistent shedding of ZIKV in these body fluids over time.

Detected ZIKV from two body fluid compartments, serum and urine, was capable of growth in cell culture from 4 of 4 inoculated animals, and thus presumably infectious. We were unable to culture virus from collected urine, fecal, and semen samples; these findings may reflect differences in relative viral loads or *bona fide* loss of infectivity in these other body fluids. The potential for ZIKV transmission through deep kissing has been raised in a recent case report describing sexual transmission of ZIKV (35). However, although ZIKV has been detected in saliva (8, 31), no cases of human transmission through saliva have been documented to date. The finding that saliva from inoculated animals, none of which developed symptoms, was infectious at 7 dpi has clinical and public health implications regarding the possibility for transmission of the virus by deep kissing or oral sex, especially in asymptomatic ZIKV-infected individuals.

Whole blood transcriptome showed maximum differential expression (1,049 DEGs) in marmosets at day 9 post-ZIKV infection. This date corresponds to the first day for which positive anti-Zika serological results were obtained, suggesting that the highest transcriptional response against ZIKV infection is triggered by antibody production.

Zika virus is sensitive to the effect of both type I and II interferons in human skin fibroblasts (36) and mice lacking type I interferon activity due to absence of the interferon-α receptor (37). Here we found by transcriptome profiling that ZIKV infection induces significant up-regulation of the type I interferon pathway at days 3, 7, and 9 post-infection (and the type II interferon pathway at day 9), consistent with an increased number of NK cells at days 7 and 9 (**Fig. 3A**) (38). However, cytokine data show increases in protein expression of type II interferons (IFN-γ and MIG) and not type I at days 3, 7, and 9 (**Fig. 3B**). ZIKV is known to inhibit the type I interferon pathway in human, (but not mouse) cells by inducing STAT2 degradation by the proteasome (14). Consistent with this finding, we observed in the current study productive ZIKV infections of marmosets and up-regulation of both type I and II interferon signaling, but only an increase in type II interferon protein expression was observed. Thus, it is possible that suppression of the type I interferon-related antiviral response in ZIKV infection in humans and non-human primates is rescued by type II interferon pathways.

Infection of human cortical neural progenitors infections showed that interferon stimulated gene MX1 was found to be specific to the infection with the Asian lineage of Zika virus, not the African lineage (39). However, although marmosets in this study were infected with the African lineage of ZIKV, they showed significant up-regulation of MX1 from day 3 to day 9. MX1 and OAS2 were also found to be prominent DEGs in ZIKV-infected human skin cells (36). These transcripts could serve as early (prior to positive serology) biomarkers of Zika infection. Small nucleolar RNA U3 is up-regulated in breast cancers and is a candidate biomarker for tumorigenesis (40). The utility of these genes as potential biomarkers for acute ZIKV infection merits further investigation.

There are several advantages to the use of marmosets as a candidate animal model for Zika infection. First, the small size of these animals relative to more common NHP models (e.g. rhesus macaques) makes them valuable in studies involving novel vaccines or therapeutics available in limited amount. Marmosets can be maintained in natural social groupings, facilitating studies of ZIKV transmission that would be difficult to accomplish with larger-bodied NHP. Finally, the results of this study and surveillance data in wild marmosets (22) reveal that these monkeys may constituting a stable reservoir for the virus in Brazil and other Latin American countries most affected by the ongoing Zika outbreak in the Americas.

## ACKNOWLEDGEMENTS

We would like to acknowledge the veterinary and pathology staff at the SNPRC for their assistance in taking care of the marmosets and collecting / analyzing samples for this study.

The animal work in this study was supported by the Southwest National Primate Research Center (P51-OD011133). This study was also supported NIH grants R01-HL105704 (CYC) and R21-AI129455 (CYC) and an award from Abbott Laboratories, Inc. (CYC). The funders had no role in study design, data collection and analysis, decision to publish, or preparation of the manuscript.

## SUPPLEMENTAL MATERIAL

**Supplementary Table 1. Scoring system used for assessing clinical symptoms in marmosets experimentally infected with ZIKV**.

**Supplementary Table 2. Detection of ZIKV and viral load quantification in longitudinally collected body fluids and necropsy tissues**.

**Supplementary Table 3. Serum levels of 22 cytokines from ZIKV-infected monkeys**.

**Supplementary Table 4. Differentially expressed genes (DEGs) shared across days 3, 7, and 9 post-infection (n=12) in ZIKV-infected marmosets**

**Supplementary Figure 1. Principal component analysis (PCA) of the gene expression profiles of ZIKV-infected marmosets and controls**. A two-dimensional PCA plot comparing ZIKV-infected marmosets (circles, color-coded by day post-inoculation) and uninfected controls (green triangles) is shown. Gene expression profiles were obtained by whole blood transcriptome analysis. No apparent clustering suggestive of technical bias is observed.

**Supplementary Figure 2. Transcriptome coverage of whole blood samples from ZIKV-infected marmosets and controls**. The bar graph shows the number of “preprocessed” reads, or reads remaining after removing low-quality (Phred score < 30) and short (length <100 bp) sequences. The line graph shows the transcriptome coverage as the percentage of gene isoforms with nonzero counts.

**Supplementary Figure 3. Type I and II interferon pathways activated during acute ZIKV infection**. Shown are transcripts associated with the type I and type II interferon pathways at days 3, 7, and 9 post-infection (red = transcript up-regulation; green = transcript down-regulation; orange = predicted activation, with darker shades of orange reflecting increased levels of activation; blue = predicted inhibition).

## REFERENCES

1. Focosi D, Maggi F, Pistello M. 2016. Zika Virus: Implications for Public Health. Clin Infect Dis 63:227–33.

2. Brasil P, Pereira JP, Jr., Moreira ME, Ribeiro Nogueira RM, Damasceno L, Wakimoto M, Rabello RS, Valderramos SG, Halai UA, Salles TS, Zin AA, Horovitz D, Daltro P, Boechat M, Raja Gabaglia C, Carvalho de Sequeira P, Pilotto JH, Medialdea-Carrera R, Cotrim da Cunha D, Abreu de Carvalho LM, Pone M, Machado Siqueira A, Calvet GA, Rodrigues Baiao AE, Neves ES, Nassar de Carvalho PR, Hasue RH, Marschik PB, Einspieler C, Janzen C, Cherry JD, Bispo de Filippis AM, Nielsen-Saines K. 2016. Zika Virus Infection in Pregnant Women in Rio de Janeiro. N Engl J Med 375:2321–2334.

3. Duffy MR, Chen TH, Hancock WT, Powers AM, Kool JL, Lanciotti RS, Pretrick M, Marfel M, Holzbauer S, Dubray C, Guillaumot L, Griggs A, Bel M, Lambert AJ, Laven J, Kosoy O, Panella A, Biggerstaff BJ, Fischer M, Hayes EB. 2009. Zika virus outbreak on Yap Island, Federated States of Micronesia. N Engl J Med 360:2536–43.

4. Carteaux G, Maquart M, Bedet A, Contou D, Brugieres P, Fourati S, Cleret de Langavant L, de Broucker T, Brun-Buisson C, Leparc-Goffart I, Mekontso Dessap A. 2016. Zika Virus Associated with Meningoencephalitis. N Engl J Med 374:1595–6.

5. Dos Santos T, Rodriguez A, Almiron M, Sanhueza A, Ramon P, de Oliveira WK, Coelho GE, Badaro R, Cortez J, Ospina M, Pimentel R, Masis R, Hernandez F, Lara B, Montoya R, Jubithana B, Melchor A, Alvarez A, Aldighieri S, Dye C, Espinal MA. 2016. Zika Virus and the Guillain-Barre Syndrome - Case Series from Seven Countries. N Engl J Med 375:1598–1601.

6. Russell K, Hills SL, Oster AM, Porse CC, Danyluk G, Cone M, Brooks R, Scotland S, Schiffman E, Fredette C, White JL, Ellingson K, Hubbard A, Cohn A, Fischer M, Mead P, Powers AM, Brooks JT. 2017. Male-to-Female Sexual Transmission of Zika Virus-United States, January-April 2016. Clin Infect Dis 64:211–213.

7. Mansuy JM, Pasquier C, Daudin M, Chapuy-Regaud S, Moinard N, Chevreau C, Izopet J, Mengelle C, Bujan L. 2016. Zika virus in semen of a patient returning from a non-epidemic area. Lancet Infect Dis 16:894–5

8. Bingham AM, Cone M, Mock V, Heberlein-Larson L, Stanek D, Blackmore C, Likos A. 2016. Comparison of Test Results for Zika Virus RNA in Urine, Serum, and Saliva Specimens from Persons with Travel-Associated Zika Virus Disease - Florida, 2016. MMWR Morb Mortal Wkly Rep 65:475–8

9. Motta IJ, Spencer BR, Cordeiro da Silva SG, Arruda MB, Dobbin JA, Gonzaga YB, Arcuri IP, Tavares RC, Atta EH, Fernandes RF, Costa DA, Ribeiro LJ, Limonte F, Higa LM, Voloch CM, Brindeiro RM, Tanuri A, Ferreira OC, Jr. 2016. Evidence for Transmission of Zika Virus by Platelet Transfusion. N Engl J Med 375:1101–3

10. Cugola FR, Fernandes IR, Russo FB, Freitas BC, Dias JL, Guimaraes KP, Benazzato C, Almeida N, Pignatari GC, Romero S, Polonio CM, Cunha I, Freitas CL, Brandao WN, Rossato C, Andrade DG, Faria Dde P, Garcez AT, Buchpigel CA, Braconi CT, Mendes E, Sall AA, Zanotto PM, Peron JP, Muotri AR, Beltrao-Braga PC. 2016. The Brazilian Zika virus strain causes birth defects in experimental models. Nature 534:267–71

11. Lazear HM, Govero J, Smith AM, Platt DJ, Fernandez E, Miner JJ, Diamond MS. 2016. A Mouse Model of Zika Virus Pathogenesis. Cell Host Microbe 19:720–30

12. Li C, Xu D, Ye Q, Hong S, Jiang Y, Liu X, Zhang N, Shi L, Qin CF, Xu Z. 2016. Zika Virus Disrupts Neural Progenitor Development and Leads to Microcephaly in Mice. Cell Stem Cell 19:672

13. Miner JJ, Cao B, Govero J, Smith AM, Fernandez E, Cabrera OH, Garber C, Noll M, Klein RS, Noguchi KK, Mysorekar IU, Diamond MS. 2016. Zika Virus Infection during Pregnancy in Mice Causes Placental Damage and Fetal Demise. Cell 165:1081–91

14. Grant A, Ponia SS, Tripathi S, Balasubramaniam V, Miorin L, Sourisseau M, Schwarz MC, Sanchez-Seco MP, Evans MJ, Best SM, Garcia-Sastre A. 2016. Zika Virus Targets Human STAT2 to Inhibit Type I Interferon Signaling. Cell Host Microbe 19:882–90

15. Dudley DM, Aliota MT, Mohr EL, Weiler AM, Lehrer-Brey G, Weisgrau KL, Mohns MS, Breitbach ME, Rasheed MN, Newman CM, Gellerup DD, Moncla LH, Post J, Schultz-Darken N, Schotzko ML, Hayes JM, Eudailey JA, Moody MA, Permar SR, O'Connor SL, Rakasz EG, Simmons HA, Capuano S, Golos TG, Osorio JE, Friedrich TC, O'Connor DH. 2016. A rhesus macaque model of Asian-lineage Zika virus infection. Nat Commun 7:12204

16. Carrion R, Jr., Patterson JL. 2012. An animal model that reflects human disease: the common marmoset (Callithrix jacchus). Curr Opin Virol 2:357–62

17. Carrion R, Jr., Ro Y, Hoosien K, Ticer A, Brasky K, de la Garza M, Mansfield K, Patterson JL. 2011. A small nonhuman primate model for filovirus-induced disease. Virology 420:117–24

18. Carrion R Jr, Brasky K, Mansfield K, Johnson C, Gonzales M, Ticer A, Lukashevich I, Tardif S, Patterson J 2007. Lassa virus infection in experimentally infected marmosets: liver pathology and immunophenotypic alterations in target tissues. J Virol 816482–90

19. Yu G, Yagi S, Carrion R Jr., Chen EC, Liu M, Brasky KM, Lanford RE, Kelly KR, Bales KL, Schnurr DP, Canfield DR, Patterson JL, Chiu CY. 2013. Experimental cross-species infection of common marmosets by titi monkey adenovirus. PLoS One 8:e68558

20. Moi ML, Ami Y, Shirai K, Lim CK, Suzaki Y, Saito Y, Kitaura K, Saijo M, Suzuki R, Kurane I, Takasaki T. 2015. Formation of infectious dengue virus-antibody immune complex in vivo in marmosets (Callithrix jacchus) after passive transfer of anti-dengue virus monoclonal antibodies and infection with dengue virus. Am J Trop Med Hyg 92:370–6

21. Verstrepen BE, Fagrouch Z, van Heteren M, Buitendijk H, Haaksma T, Beenhakker N, Palu G, Richner JM, Diamond MS, Bogers WM, Barzon L, Chabierski S, Ulbert S, Kondova I, Verschoor EJ. 2014. Experimental infection of rhesus macaques and common marmosets with a European strain of West Nile virus. PLoS Negl Trop Dis 8:e2797

22. Favoretto S, Araujo D, Oliveira D, Duarte N, Mesquita F, Zanotto P, Durigon E. 2016. First detection of Zika virus in neotropical primates in Brazil a possible new reservoir. bioRxiv doi:https://doi.org/10.1101/049395.

23. Dick GW, 1953. Epidemiological notes on some viruses isolated in Uganda; Yellow fever, Rift Valley fever, Bwamba fever, West Nile, Mengo, Semliki forest, Bunyamwera, Ntaya, Uganda S and Zika viruses. Trans R Soc Trop Med Hyg 47:13–48

24. Lanciotti RS, Kosoy OL, Laven JJ, Velez JO, Lambert AJ, Johnson AJ, Stanfield SM, Duffy MR. 2008. Genetic and serologic properties of Zika virus associated with an epidemic, Yap State, Micronesia, 2007. Emerg Infect Dis 14:1232–9

25. Rabe IB, Staples JE, Villanueva J, Hummel KB, Johnson JA, Rose L, Mts, Hills S, Wasley A, Fischer M, Powers AM. 2016. Interim Guidance for Interpretation of Zika Virus Antibody Test Results. MMWR Morb Mortal Wkly Rep 65:543–6

26. Giavedoni LD, 2005. Simultaneous detection of multiple cytokines and chemokines from nonhuman primates using luminex technology. J Immunol Methods 301:89–101

27. Dobin A, Davis CA, Schlesinger F, Drenkow J, Zaleski C, Jha S, Batut P, Chaisson M, Gingeras TR. 2013. STAR: ultrafast universal RNA-seq aligner. Bioinformatics 29:15–21

28. Anders S, Pyl PT, Huber W. 2015. HTSeq–a Python framework to work with high-throughput sequencing data. Bioinformatics 31:166–9

29. Robinson MD, McCarthy DJ, Smyth GK. 2010. edgeR: a Bioconductor package for differential expression analysis of digital gene expression data Bioinformatics 26:139–40

30. Corman VM, Rasche A, Baronti C, Aldabbagh S, Cadar D, Reusken CB, Pas SD, Goorhuis A, Schinkel J, Molenkamp R, Kummerer BM, Bleicker T, Brunink S, Eschbach-Bludau M, Eis-Hubinger AM, Koopmans MP, Schmidt-Chanasit J, Grobusch MP, de Lamballerie X, Drosten C, Drexler JF. 2016. Assay optimization for molecular detection of Zika virus. Bull World Health Organ 94:880–892

31. Barzon L, Pacenti M, Berto A, Sinigaglia A, Franchin E, Lavezzo E, Brugnaro P, Palu G. 2016. Isolation of infectious Zika virus from saliva and prolonged viral RNA shedding in a traveller returning from the Dominican Republic to Italy, January 2016 Euro Surveill 21

32. Barzon L, Pacenti M, Franchin E, Lavezzo E, Trevisan M, Sgarabotto D, Palu G. 2016. Infection dynamics in a traveller with persistent shedding of Zika virus RNA in semen for six months after returning from Haiti to Italy, January 2016 Euro Surveill 21

33. Lorenzo-Redondo R, Fryer HR, Bedford T, Kim EY, Archer J, Kosakovsky Pond SL, Chung YS, Penugonda S, Chipman JG, Fletcher CV, Schacker TW, Malim MH, Rambaut A, Haase AT, McLean AR, Wolinsky SM. 2016. Persistent HIV-1 replication maintains the tissue reservoir during therapy. Nature 530:51–6

34. Weitzel T, Cortes CP. 2016. Zika Virus Infection Presenting with Postauricular Lymphadenopathy. Am J Trop Med Hyg 95:255–6

35. D'Ortenzio E, Matheron S, Yazdanpanah Y, de Lamballerie X, Hubert B, Piorkowski G, Maquart M, Descamps D, Damond F, Leparc-Goffart I. 2016. Evidence of Sexual Transmission of Zika Virus. N Engl J Med 374:2195–8

36. Hamel R, Dejarnac O, Wichit S, Ekchariyawat P, Neyret A, Luplertlop N, Perera-Lecoin M, Surasombatpattana P, Talignani L, Thomas F, Cao-Lormeau VM, Choumet V, Briant L, Despres P, Amara A, Yssel H, Misse D. 2015. Biology of Zika Virus Infection in Human Skin Cells. J Virol 89:8880–96

37. Rossi SL, Tesh RB, Azar SR, Muruato AE, Hanley KA, Auguste AJ, Langsjoen RM, Paessler S, Vasilakis N, Weaver SC. 2016. Characterization of a Novel Murine Model to Study Zika Virus. Am J Trop Med Hyg 94:1362–9

38. Guan J, Miah SM, Wilson ZS, Erick TK, Banh C, Brossay L. 2014. Role of type I interferon receptor signaling on NK cell development and functions. PLoS One 9:e111302

39. Zhang F, Hammack C, Ogden SC, Cheng Y, Lee EM, Wen Z, Qian X, Nguyen HN, Li Y, Yao B, Xu M, Xu T, Chen L, Wang Z, Feng H, Huang WK, Yoon KJ, Shan C, Huang L, Qin Z, Christian KM, Shi PY, Xu M, Xia M, Zheng W, Wu H, Song H, Tang H, Ming GL, Jin P. 2016. Molecular signatures associated with ZIKV exposure in human cortical neural progenitors. Nucleic Acids Res 44:8610–8620

40. Langhendries JL, Nicolas E, Doumont G, Goldman S, Lafontaine DL. 2016. The human box C/D snoRNAs U3 and U8 are required for pre-rRNA processing and tumorigenesis. Oncotarget 7:59519–59534

